# Age-related differences in metabolome composition in a healthy pediatric population: A pilot analysis

**DOI:** 10.64898/2026.09.13.750246

**Authors:** Amy Thachil, Rupasri Mandal, David S. Wishart, Tom Blydt-Hansen

**Affiliations:** University of British Columbia, Department of Pediatrics, Vancouver, British Columbia, Canada; University of Alberta, Department of Biological Sciences, Edmonton, Alberta, Canada; University of Alberta, Department of Computing Sciences, Edmonton, Alberta, Canada; University of Alberta, Department of Pathology and Laboratory Medicine, Edmonton, Alberta, Canada

**Keywords:** Metabolomics, age, pediatrics, biomarkers, growth, development, puberty, metabolism

## Abstract

Metabolic profiling of biofluids provides a comprehensive overview of physiological processes occurring at the molecular, cellular, tissue, and organ levels. Accounting for age-related variation in the metabolome is critical to accurately interpret these effects in blood-based metabolomics research. In this study, we sought to characterize the blood metabolome of healthy pediatric patients and assess whether metabolite concentrations differ across age groups and stages of sexual maturation. A total of 44 plasma samples from individuals aged 0–19 years were analyzed using a targeted quantitative metabolomics approach, combining direct injection with reverse-phase liquid chromatography mass spectrometry. Samples were stratified by age: infants (0–2 years), pre-teens (10–12 years), and post-pubertal adolescents (16–19 years). Asymmetric dimethylarginine, carnosine, N-acetyl-serine, and quinolinic acid had a negative association with age, whereas creatinine, homoarginine, lysoPC a C16:1, guanidoacetic acid and 5-Hydroxyindoleacetic acid had a positive association with age. Comparisons between the pre-teen and infant age groups yielded the greatest number of differences followed by the post-pubertal versus infant and post-pubertal versus pre-teen groups. These findings demonstrate age-associated heterogeneity in the pediatric plasma metabolome. Establishing normative reference ranges for metabolite concentrations will be essential to improve the accuracy of metabolomic analyses that include pediatric participants.

## Introduction

Metabolic profiling of biofluids and tissue samples provides a snapshot of cellular metabolism and the physiological mechanisms occurring at a molecular, cellular, tissue and organ level (1). Differences in blood metabolome composition can be attributed to environmental and genetic influences on physiology. Biological sex, immune status, nutritional state, and disease state are amongst the innate host factors that may impact metabolome composition (2–4). Given the changes in physiology and metabolism that accompany human development, it is to be expected that metabolome composition varies with age. Accounting for age-related variation is essential for accurate interpretation in blood metabolomics research, particularly in biomarker discovery.

Age-related changes in blood metabolome composition are well described within the adult population. Saito et al. (5) identified age and sex-related differences in serum metabolome composition in 60 adult Japanese participants categorized as: young males (25–35 years old), older males (55–64 years old), young females (25–35 years old), and older females (55–65 years old). 130 out of 516 serum metabolites were found to be significantly different between the young and old females (p<0.05), whereas 158 metabolites were found to differ according to age in males. These findings were further corroborated by additional reports from Yu et al. (6), the Husermet project (7) and Frederiksen et al (8). Yu et al. found that serum concentrations of: C0, C10:1, C12:1, C18:1, SM C16:1, SM C18:1, and PC aa C28:1 increased, while histidine decreased with age in adult female patients, after controlling for BMI (6). The Husermet project also identified age-related differences in the human serum metabolome, in addition to differences associated with sex, BMI, blood pressure and smoking (7). Work by Frederiksen et al (8). identified sex and age specific reference ranges for 16 steroid metabolites including: progesterone, 17-hydroxypregnenolone, 17-hydroxyprogesterone, dehydroepiandrosterone sulfate, androstenedione, testosterone, dihydrotestosterone, 11-deoxycorticosterone, corticosterone, 11-deoxycortisol, cortisol, and cortisone (n=2458, age ranges: 0-77 years). Together, these studies demonstrate that age is a consistent determinant of metabolite profiles across adult populations, underscoring the need to account for age as a covariate in metabolomics analyses. This also highlights the importance of extending such investigations to earlier stages of life where the metabolome is likely shaped by rapid growth and development.

Age-related differences in the metabolome composition have been reported in children, but to a lesser extent. One group conducted a large meta-analysis of 6 separate European pediatric cohorts consisting of 1192 children total, aged 6–11 (9). Of the 177 serum metabolites and 44 urinary metabolites measured, only serum and urinary creatinine levels were found to be positively associated with age, across all cohorts. Another analysis conducted by Hirschel et al. (10), examined 23 amino acid and 6 acylcarnitine levels in dried whole blood samples obtained from 191 patients aged 3 months to 18 years. They identified 4 distinct trends in metabolite concentration within this age cohort: 1) decreasing concentration following infancy and reaching stable concentrations by 1-5 years of age, 2) a continuous increase in concentration from the age of 3 months to 18 years, 3) concentration increase until 5-9 years of age and 4) no discernible trend in concentration change with age. Metabolites were found to follow heterogeneous and metabolite-specific trajectories, underscoring the need for age-stratified reference ranges and longitudinal studies to accurately characterize developmental metabolic patterns in children.

In contrast to adults, the pediatric metabolome is expected to be highly dynamic and closely linked to developmental milestones. Hormonal shifts, such as changes in growth hormone (11, 12), insulin-like growth factor (13, 14), and sex hormones (15), play a major role in shaping metabolic pathways, including those involved in amino acid, lipid, and energy metabolism. Nutritional intake (16–18) and the gut microbiome (19, 20) also change substantially from infancy through adolescence, which can influence metabolite profiles. Sex differences in certain metabolites, such as sphingolipids, have been reported as early as infancy (21) and into early childhood (22), and may become more pronounced during and after pubertal development (23, 24). These observations highlight that the pediatric metabolome maybe shaped by coordinated physiological changes, including growth, hormonal maturation, and nutrition. There is need for tailored frameworks when interpreting metabolomic data in children.

Previous analyses have not adequately captured metabolomic differences across key developmental stages, including infancy, adolescence, and the post-pubertal period. Such characterization is essential to advance our understanding of metabolic changes across childhood and to enable appropriate adjustment for age- and development-related variation in pediatric metabolomics studies. Here, we aim to characterize the blood metabolome of healthy children to explore whether there are differences in metabolite concentrations across various ages and stages of sexual maturation. This pilot analysis will inform a larger study characterizing metabolome changes throughout childhood development, and establishing references ranges for blood metabolite concentrations across the pediatric age spectrum.

## Methods

### Sample identification

We conducted a retrospective, observational pediatric cohort study using plasma samples from children enrolled in the institutional biobank at a single tertiary pediatric hospital. As part of the biobank program, deidentified blood samples are collected from individuals attending different clinical areas, often concurrently with routine procedures. Our aim was to identify approximately 50 plasma samples from children at different stages of sexual maturation. Samples were considered for inclusion and stratified based on patient age at the time of sample retrieval: infants (0-2 years of age), pre-teens (10-12 years of age) and post-pubertal (16-19 years of age). The age groups presented here were selected with the consideration that development-related changes in metabolome composition would be most pronounced when comparing individuals at palpably different points of development: infancy, pre-puberty, and the later teenage years. The pre-teen and post-pubertal groups are closer to adulthood, making it reasonable that less variation would exist between these stages.

Patients were excluded to avoid measuring inflammation-related artefacts in metabolome composition based on reasons for hospitalization and primary diagnosis. Based on this premise, samples obtained from recipients of solid organ transplants and patients with acute infections, systemic inflammatory conditions and/or metabolic disorders were not included in the analysis. The institutional biobank with patient consent was approved by the local research ethics board (REB # H13-0311), and separate approval was obtained for this study, without requirement for additional consent (REB # H22-02845).

Patient information was obtained from institutional biobank records at the time of sample retrieval. Datapoints available for analysis included: patient age, sex, referral department, diagnosis, and sample storage details.

### Liquid chromatography with tandem mass spectrometry metabolomics testing

Blood metabolomic analysis was conducted at The Metabolomic Innovation Centre (TMIC, Edmonton) using a targeted quantitative metabolomics approach to analyze samples with a combination of direct injection mass spectrometry (DI-MS) and reverse phase LC-MS/MS. The TMIC MEGA panel was used in conjunction with the SCIEX Qtrap® 5500 tandem mass spectrometry instrument (AB SCIEX, Framingham, MA) equipped with a solvent delivery system to quantify 635 unique plasma metabolites in each sample tested. All samples were processed according to established protocols for the MEGA kit and mass spectrometric apparatus as described previously (25). Metabolomic data analysis and concentration calculations were obtained using the SCIEX Analyst software.

### Data cleaning

Missing concentration values were imputed with the assay limit of detection divided by the square root of 2. Metabolites with over 75% missing values were removed, leaving 566 metabolites for analysis. Metabolite concentrations were log10 transformed prior to performing statistical analysis. A final dataset was prepared merging metabolite concentration data and available patient clinical characteristics and imported into the R Studio environment for statistical analysis and the MetaboAnalyst 5.0 software (26) for visualization.

### Statistical analysis

Metabolite data were visualized using the principal component analysis (PCA) and the MetaboAnalyst 5.0 software (26). For the purposes of PCA, missing values were replaced by the assay limit of detection, metabolites with over 75% missing values were removed, and concentrations were log10 transformed. The data were also “auto-scaled,”; metabolite concentrations were mean-centred and expressed in standard deviation units.

Metabolite concentrations were visualized using volcano plots to explore the relationships between age groups. Comparisons were made between the following group pairings: pre-teen/infant, post-pubertal/pre-teen and post-pubertal/infant age groups. Raw metabolite concentrations were used to calculate the fold changes in concentration between age groups. Fold changes were reported in reference to the older age group and were log2 transformed prior to being plotted.

Tests for association were first evaluated using a significance level of p <0.05 and then after Bonferroni correction to limit the risk of introducing Type I error with multiple comparisons (p <9.0×10^-5^). Univariate correlations between age and individual metabolite concentration were determined using a Spearman’s rank test to account for the non-normal distribution of concentrations. To evaluate group-related differences in metabolome composition between the infant, pre-teen and post-pubertal stages, we conducted an analysis of variance (ANOVA) test for each metabolite. All metabolites demonstrating a difference in group mean according to ANOVA were subsequently evaluated using the Tukey-Kramer test.

The small size of our cohort limited our ability to assess sex differences within age groups. As such, sex differences were assessed at the level of the whole cohort. For each metabolite, a two-sample t-test was used to determine whether the mean concentration differed between males and females.

## Results

### Patient population and sample selection

48 plasma samples from patients between the ages of 0 and 19 were available for metabolomic testing. 4 samples obtained from individuals diagnosed with viral encephalitis, cystitis and encephalopathy were excluded on the basis of not meeting inclusion diagnosis criteria. The 44 plasma samples included in the analysis were obtained from individuals diagnosed with allergies, spinal muscular atrophy, epilepsy, tonsilitis, sleep apnea, and hypertrophy of the adenoids. The infant group consisted of 18 samples, the pre-teen group of 17 samples and the post-pubertal group of 9 samples. A breakdown of available cohort characteristics can be found in **Table 1**.

**Table 1.** Patient baseline characteristics at the time of sample retrieval. Values are expressed as a count (percentage of a group).

|  | All patients (n=44) |
| --- | --- |
| **Age** |  |
| 0-2 | 18 (41) |
| 10-12 | 17 (39) |
| 16-19 | 9 (20) |
| **Female Sex** | 18 (41) |
| 0-2 | 8 (44) |
| 10-12 | 7 (41) |
| 16-19 | 3 (33) |
| **Diagnosis Department** |  |
| Allergy | 30 (68) |
| Ear, Nose and Throat (ENT) | 5 (12) |
| Neurology | 9 (20) |

Patients included in the analysis were overwhelmingly referred by the allergy department, although select samples were also obtained from patients in the ear, nose and throat (ENT) and neurology departments. The infant and pre-teen age cohorts had similar proportions of males and females; 44% of the infant samples were obtained from females and 41% of the pre-teen samples were obtained from females. The post-pubertal group was male-dominant, with only 33% of samples obtained from females.

### Age and metabolite concentration correlations

143 metabolites demonstrated a significant univariate association with age at the time of sample retrieval (p<0.05). 9 metabolites retained a significant association after accounting for multiple comparisons (**Table 2**) (p<9.0×10^-5^); Asymmetric dimethylarginine, carnosine, N-Acetyl-Serine, and quinolinic acid had a negative association with age, whereas creatinine, homoarginine, lysoPC a C16:1, guanidoacetic acid and 5-Hydroxyindoleacetic acid had a positive association with age. Smaller p-values were observed for metabolites showing stronger correlation with age, as measured by absolute Spearman’s rho value. (**Figure 1**).

**Figure 1.**
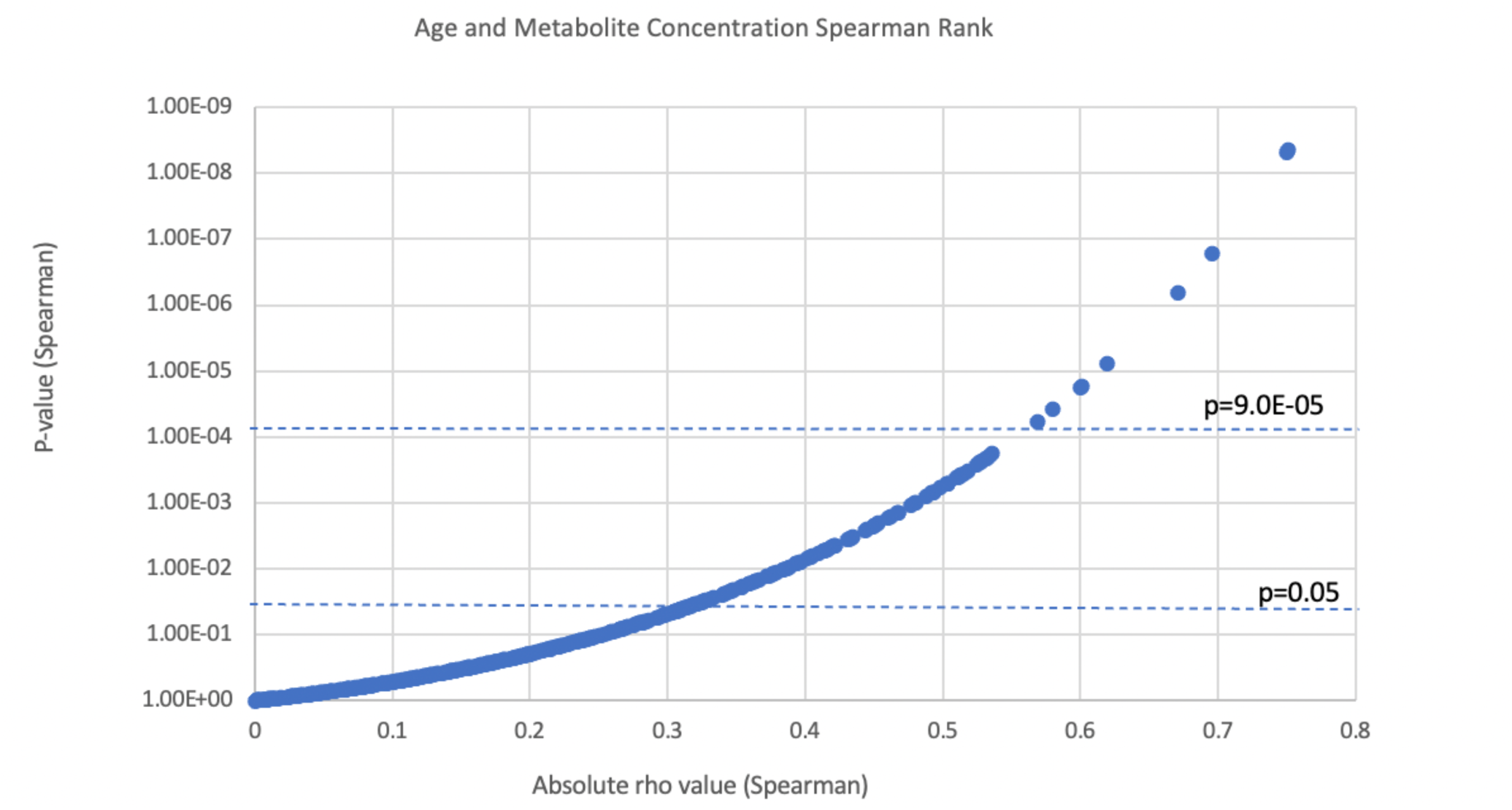
Association between absolute Spearman’s rho and p-value. The correlation between age and concentration was computed for each metabolite in the final dataset (n=566).Metabolites with a stronger univariate association were found to have increasing significance. 9 metabolites retained a significant association following adjustment for multiple comparisons.

**Table 2.** Age associated metabolites post-correction. All values rounded to the second decimal place.

| Metabolite | Spearman's rho | P-value |
| --- | --- | --- |
| Asymmetric dimethylarginine | -0.60 | 1.65E-05 |
| Carnosine | -0.57 | 5.70E-05 |
| Creatinine | 0.70 | 1.59E-07 |
| Homoarginine | 0.60 | 1.72E-05 |
| LysoPC a C16:1 | 0.62 | 7.63E-06 |
| Guanidoacetic acid | 0.75 | 4.77E-09 |
| N-Acetyl-Serine | -0.67 | 6.35E-07 |
| 5-Hydroxyindoleacetic acid | 0.58 | 3.80E-05 |
| Quinolinic acid | -0.75 | 4.36E-09 |

### Data visualization using principal component analysis and a volcano plot

All 566 metabolites were included in PCA data visualization. Visually, samples showed separation by age group along the first and second principal components (**Figure 2**). 30.5% of sample variation was accounted for by the first principal component, followed by 10.3% in the second component and 7.8% in the third component (**Figure 3**). Greater overlap was observed between samples in the pre-teen and post-pubertal age groups compared to samples from the infant age group (**Figure 2**). Samples from the infant age groups appeared to separate from the other age groups along the first three principal components (**Figure 2b**), suggesting that differences in metabolome composition exist between the infant vs. pre-teen and infant vs. post-pubertal stages. Given the overall small sample size and relative lack of samples in the post-pubertal group, further study is required to confirm this.

**Figure 2.**
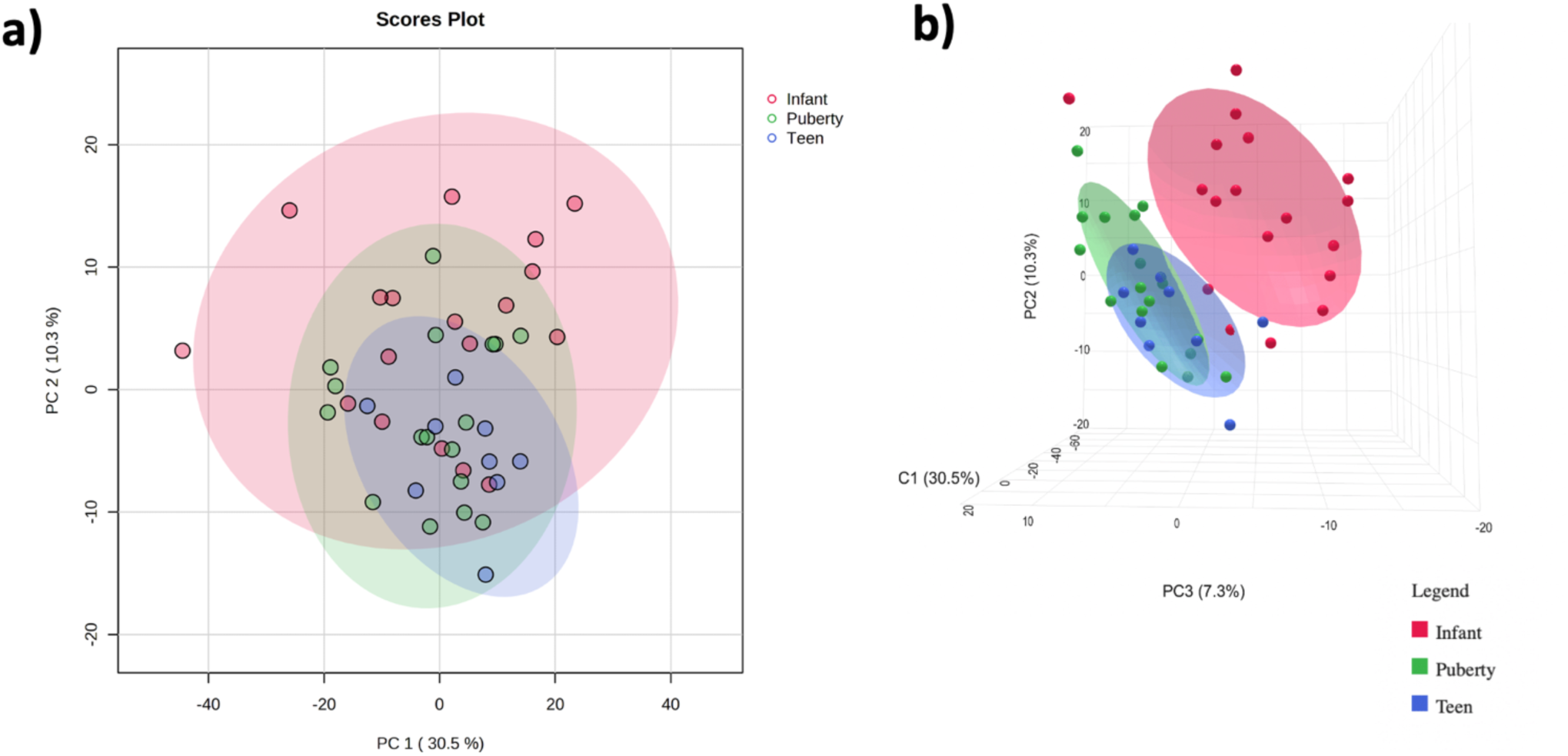
2D and 3D PCA score plots. Samples are labelled according to developmental stage: infant, pre-teen and post-pubertal. **a)** Mild sample clustering according to developmental stage is apparent in the first two components. In particular, the infant samples appear distinct from pre-teen and post-puberty samples. **b)** Infant sample clustering is apparent considering the first three components. Greater overlap is observed between pre-teen and post-puberty groups.

**Figure 3.**
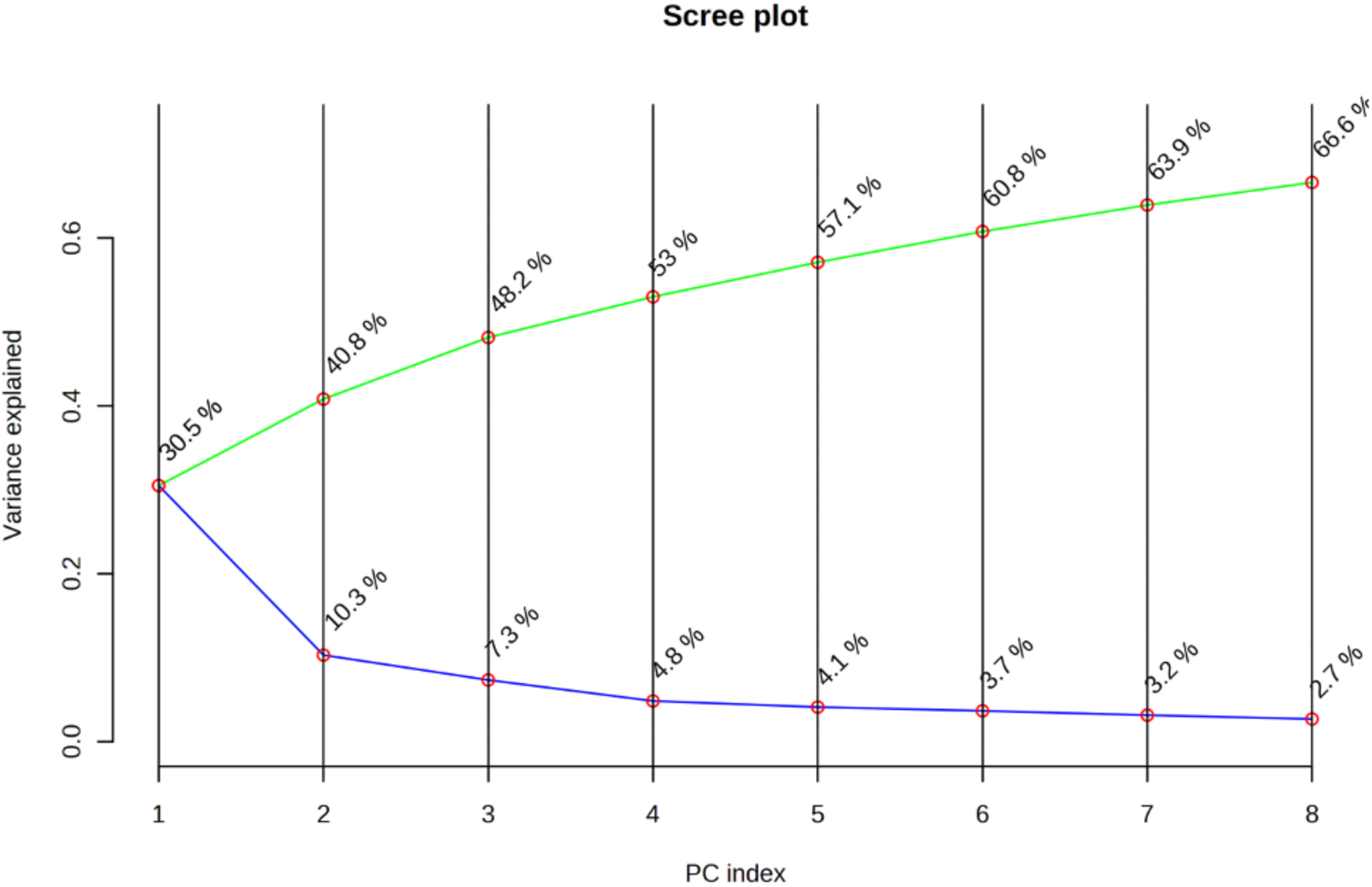
Variation explained by individual PCA components. Just under 50% of the sample variation is explained by the first 3 components of the model.

Fold change differences in metabolite concentrations between the pre-teen/infant, post-pubertal/pre-teen and post-pubertal/infant age groups are visualized in supplementary material (**Figures S1A-C).**

### Identification of metabolite differences between age groups

A series of ANOVA tests identified 115 out of the 566 metabolites that demonstrated a significant difference in mean concentration between the 3 age groups (p<0.05), of which 15 metabolites retained a significant difference in mean concentrations after accounting for multiple comparisons (p<9.0×10^-5^). These metabolites included: 5-Hydroxylysine, asymmetric dimethylarginine, carnosine, creatine, creatinine, lysine, lysoPC a C16:1, lysoPC a C17:0, PC aa C34:3, CE(18:3), 2-Hydroxyisobutyric acid, guanidoacetic acid, N-Acetyl-Serine, 5-Hydroxyindoleacetic acid, and quinolinic acid (**Table 3**) (p<9.0×10^-5^). With the exception of homoarginine, 8 out of the 9 metabolites identified as demonstrating a significant correlation with age (**Table 2**) were also among the 15 metabolites shown to differ between the three age cohorts (p<9.0×10^-5^).

**Table 3.**
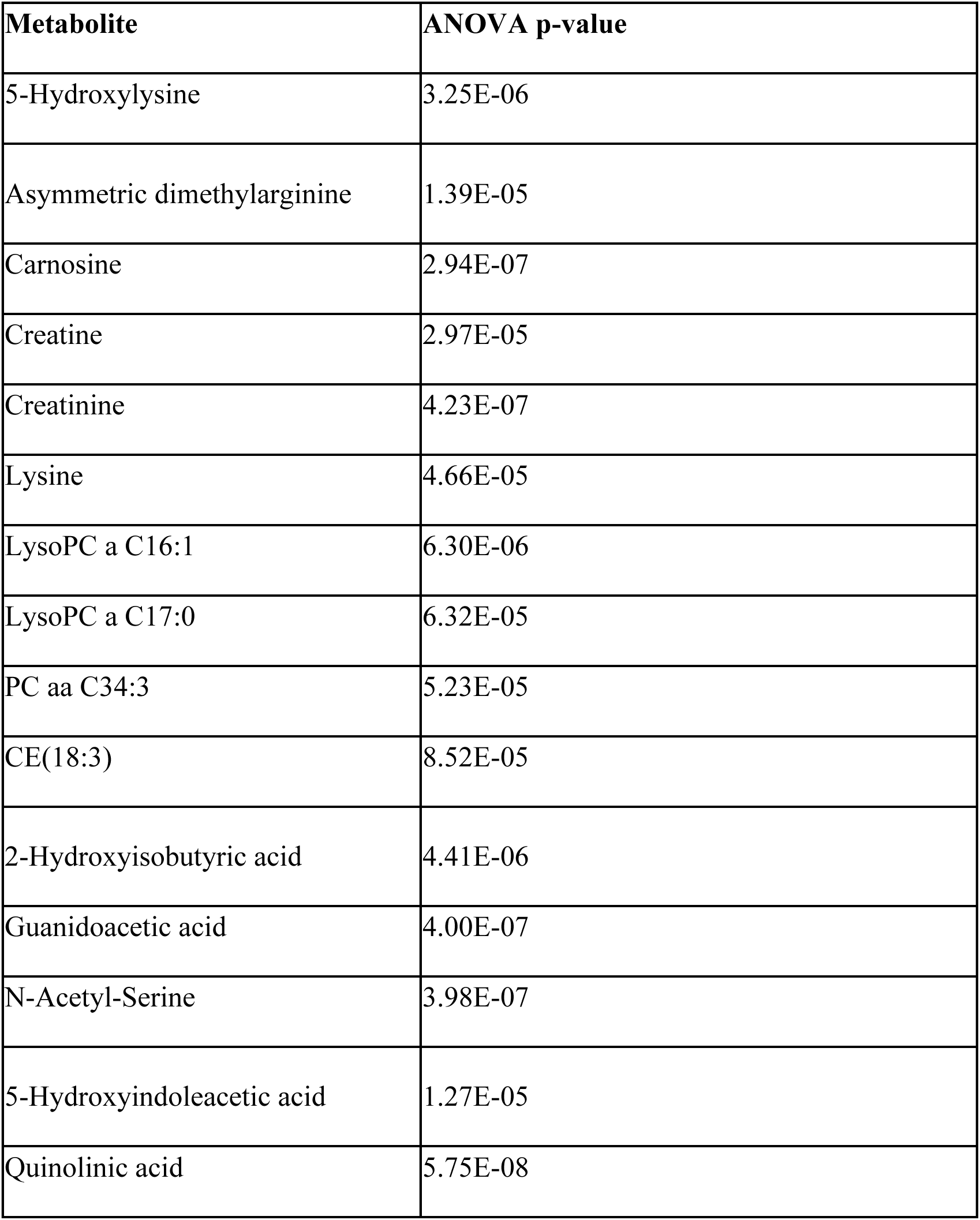
ANOVA results: Metabolites differing by developmental stage post-correction. All values are expressed to the second decimal place. Metabolites are presented according to order in assay panel.

All initial 115 metabolites demonstrating a difference in group mean concentration (p<0.05) were subsequently evaluated using a Tukey-Kramer test to identify specific group differences. Greater overlap was observed between the pre-teen and post-pubertal groups compared to the infant group; more metabolites were significantly different when comparing the post-pubertal/infant and pre-teen/infant groups versus the post-pubertal/pre-teen groups. Significant differences (p<0.05) in mean concentration between the infant and pre-teen age groups were identified in 75 metabolites; 11 metabolites between the pre-teen and post-pubertal groups; and 67 metabolites between the infant and post-pubertal groups.

Using a Bonferroni-adjusted p-value (p<9.0×10^-5^), 8 metabolites demonstrated a significant difference in mean concentration between the infant and pre-teen age groups, 1 metabolite demonstrated a significant difference between the pre-teen and post-pubertal groups, and 8 metabolites demonstrated a significant difference between the infant and post-pubertal groups (**Table 4**). 6 unique metabolites were uniquely different between the infant and pre-teen age groups, whereas 5 were uniquely different between the infant and post-pubertal groups. Two metabolites (Carnosine and N-Acetyl-Serine) were significantly higher in infants, compared with both pre-teen and post-pubertal age groups. 5-Hydroxylysine was significantly lower in the post-pubertal group, when comparing the post-pubertal/infant and post-pubertal/pre-teen age groups.

**Table 4.**
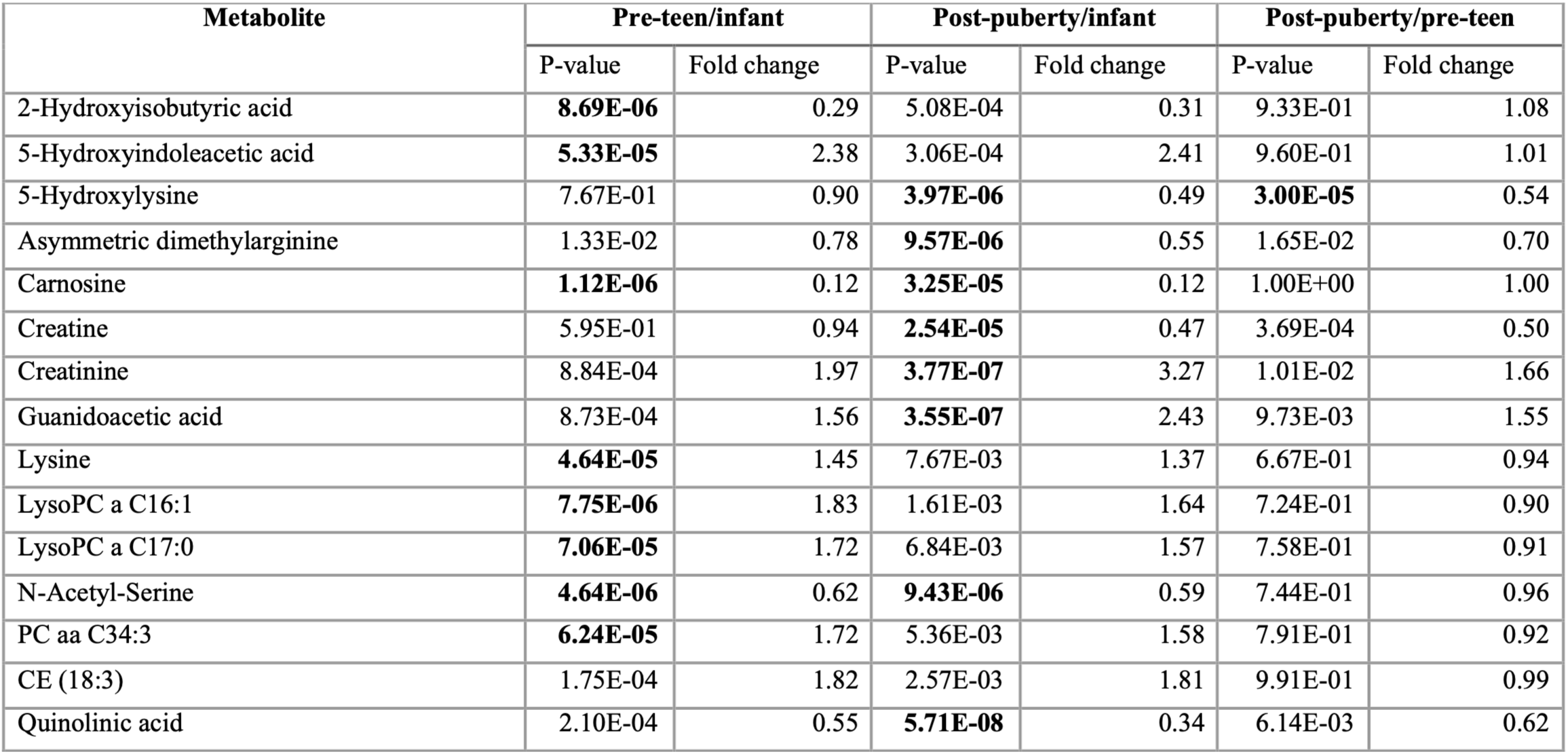
Tukey Kramer results: Comparison of metabolite concentrations between developmental stages. Fold changes were calculated in the following directions: pre-teen/infant, post-puberty/infant, post-puberty/pre-teen. A Bonferroni-corrected significance threshold of p=9.0×10^-5^ was used to identify significant differences in concentration between age groups.

| Metabolite | Pre-teen/infant |  | Post-puberty/infant |  | Post-puberty/pre-teen |  |
| --- | --- | --- | --- | --- | --- | --- |
|  | P-value | Fold change | P-value | Fold change | P-value | Fold change |
| 2-Hydroxyisobutyric acid | **8.69E-06** | 0.29 | 5.08E-04 | 0.31 | 9.33E-01 | 1.08 |
| 5-Hydroxyindoleacetic acid | **5.33E-05** | 2.38 | 3.06E-04 | 2.41 | 9.60E-01 | 1.01 |
| 5-Hydroxylysine | 7.67E-01 | 0.90 | **3.97E-06** | 0.49 | **3.00E-05** | 0.54 |
| Asymmetric dimethylarginine | 1.33E-02 | 0.78 | **9.57E-06** | 0.55 | 1.65E-02 | 0.70 |
| Carnosine | **1.12E-06** | 0.12 | **3.25E-05** | 0.12 | 1.00E+00 | 1.00 |
| Creatine | 5.95E-01 | 0.94 | **2.54E-05** | 0.47 | 3.69E-04 | 0.50 |
| Creatinine | 8.84E-04 | 1.97 | **3.77E-07** | 3.27 | 1.01E-02 | 1.66 |
| Guanidoacetic acid | 8.73E-04 | 1.56 | **3.55E-07** | 2.43 | 9.73E-03 | 1.55 |
| Lysine | **4.64E-05** | 1.45 | 7.67E-03 | 1.37 | 6.67E-01 | 0.94 |
| LysoPC a C16:1 | **7.75E-06** | 1.83 | 1.61E-03 | 1.64 | 7.24E-01 | 0.90 |
| LysoPC a C17:0 | **7.06E-05** | 1.72 | 6.84E-03 | 1.57 | 7.58E-01 | 0.91 |
| N-Acetyl-Serine | **4.64E-06** | 0.62 | **9.43E-06** | 0.59 | 7.44E-01 | 0.96 |
| PC aa C34:3 | **6.24E-05** | 1.72 | 5.36E-03 | 1.58 | 7.91E-01 | 0.92 |
| CE (18:3) | 1.75E-04 | 1.82 | 2.57E-03 | 1.81 | 9.91E-01 | 0.99 |
| Quinolinic acid | 2.10E-04 | 0.55 | **5.71E-08** | 0.34 | 6.14E-03 | 0.62 |

### Assessment of sex differences

Minimal sex differences were observed in metabolite concentrations. 16 metabolites, including carnosine, urea, glycine, and creatinine demonstrated some degree of difference between sexes (p<0.05). No metabolites demonstrated significant sex differences after correction for multiple comparisons. In the post-pubertal group (n=9; 33% female) where sex differences are most likely to be identified, the sample was too small for meaningful statistical comparison.

## Discussion

In this study, we observed marked differences in plasma metabolome composition across childhood developmental stages. Over 100 of the evaluated plasma metabolites were found to differ according to age or developmental stage, with select metabolites retaining pronounced concentration differences even after accounting for multiple comparisons. Peptides, amino acids, carboxylic acids, and lipids were among the metabolite classes found to differ. We did not identify sex-related concentration differences in the whole cohort. Based on unsupervised clustering and group-wise comparisons, we report greater differences in metabolome composition between infants and pre-teens and infants and post-pubertal teenagers compared to between the pre-teen and post-pubertal stages. A number of metabolite concentrations increased with age, whilst others were found to decrease.

To contextualize our findings, we consulted the existing literature to evaluate our results in relation to current knowledge. The literature on pediatric age-related differences in metabolome composition is limited, particularly when considering blood metabolomics work. Given these limitations, we also examined metabolite reference intervals reported in the Human Metabolome Database (HMDB) and the Canadian Laboratory Initiative on Pediatric Reference Intervals (CALIPER) databases. The HMDB is an electronic resource housing chemical, clinical and molecular-level information on metabolites (27). The CALIPER database is a pediatric-specific database providing reference levels for over 200 blood analytes (28).

Several peptide-based and carboxylic acid metabolites demonstrated age-associated variation across the pediatric cohorts examined in this study, although comprehensive pediatric reference intervals for these analytes are not yet established. Asymmetric dimethylarginine decreased with age, with lower concentrations in pre-teens and the post-pubertal group compared to infants. According to the HMDB, this compound is a by-product of protein modification, with reported pediatric blood values ranging from 0.595–1.03 µM in children between the ages of 1–13 years (29). However, previous data do not reflect developmental trajectories, limiting direct comparison with our findings. Guanidoacetic acid and 5-hydroxyindoleacetic acid also increased with age, particularly between infants and pre-teens, while quinolinic acid decreased, most notably between infants and post-pubertal children. 2-hydroxyisobutyric acid was lowest in pre-teens, with post-pubertal values intermediate between pre-teens and infants. At the time of evaluation, HMDB indicates that pediatric blood concentration data are either unavailable or limited to disease contexts (30–32) for these metabolites, with no established pediatric reference ranges (33–36).

Amino acids and related derivatives also exhibited age-dependent patterns in our analysis; however, as with the metabolites discussed above, the existing literature is limited and age-stratified pediatric reference ranges are currently unavailable. N-acetyl-serine decreased with age, with both pre-teens and post-pubertal children showing lower levels than infants; at the time of evaluation HMDB reported no prior pediatric blood concentration studies (37). Carnosine, a dipeptide synthesized from β-alanine and histidine (38), similarly lacks pediatric reference ranges, with existing values derived only from adults, including those with Alzheimer’s disease (39). Homoarginine was found to lack reference intervals in both HMDB and CALIPER but increased with age across the pediatric groups included in our study. For 5-hydroxylysine, the HMDB suggests that newborns, infants, and children up to 18 years have blood concentration levels ranging from 0–2.5 µM, with adult values ranging from 1.8–3.2 µM (40). In contrast, our data showed lower concentrations in the older, post-pubertal group compared with infants and pre-teens. Lysine concentrations were significantly higher in pre-teens than in infants, aligning with prior reports of increasing lysine from infancy into childhood and adolescence (newborns: 71–272 µM; infants: 37–228 µM; children 6–18 years: 108–270 µM) (41, 42).

Lipids and related metabolite species further supported age-related metabolic shifts. LysoPC a C16:1 and LysoPC a C17:0 increased from infancy to pre-adolescence, with no significant change thereafter, and PC aa C34:3 was highest in pre-teens, particularly relative to infants. Cholesteryl ester CE(18:3) also trended higher in pre-teens and post-pubertal children than in infants, though not significantly after multiple-testing correction. No pediatric reference concentration data were identified for these metabolites. Together, these findings indicate age-dependent changes across various metabolite classes, while also highlighting the limited availability of normative pediatric reference ranges and underscoring the need for larger, pediatric-focused studies.

We report decreasing plasma creatine and increasing creatinine levels in children between 0 and 19 years. Pair-wise comparisons between groups revealed that creatine demonstrated the greatest difference in plasma concentration when comparing the post-pubertal group to pre-teens and infants. Creatinine was most different when comparing the infant group to pre-teens and the post-pubertal group. The difference in plasma creatinine between pre-teens and the post-pubertal group was less pronounced. Creatinine is a downstream waste-product of creatine produced through muscle metabolism and is predominantly filtered and excreted through the kidney (43). Creatine is a metabolite produced in the liver and kidney to permit on-demand muscular energy-uptake (43, 44). Previous studies have reported age-related increases in creatinine concentrations during childhood development (9, 45, 46), likely reflecting the increase in muscle mass during childhood growth. Our creatinine findings are consistent with this literature and with CALIPER reference intervals, which rise from 9–32 µM in children aged 15 days–2 years to 43–96 µM in adolescents aged 15–19 years (28, 47). The inverse association observed for plasma creatine is also consistent with prior studies reporting elevated creatine concentrations in early childhood that progressively decline with age (48–50). HMDB data indicates high creatine concentrations in newborns (approximately 84.0 ± 23.2 µM) and a wide pediatric range of 5–82 µM in children aged 1–13 years (51).

Our observations of a distinctive infant plasma metabolomic profile compared to pre-teens and older teenagers may reflect the rapid physiological and metabolic changes that occur during infancy. Blood growth hormone levels are at their highest during the neonatal stage, decrease until 6 months of age, and remain constant thereafter until pre-puberty (52–54). Another important growth factor, insulin-like growth factor 1(IGF-1), progressively increases in concentration during early childhood and into puberty (55–57). These consequential hormonal transitions relate closely to growth and metabolism (53), suggesting that a distinct infant metabolic profile is to be expected.

The developing infant microbiome may also contribute to the metabolite trends observed. The infant gut microbiota is characterized by low levels of diversity which increase following introduction to solid foods and other environmental exposures (58). Approximately 2-3 years after birth, the infant gut microbiome undergoes a gradual transition from *Lactobacillus* and *Bifidobacterium* colonization to being predominated by *Firmicutes* and *Bacteroidetes* (58, 59). Following this transition, the gut microbiome remains relatively stable into adulthood (58). The interplay between the gut microbiome and metabolism influences vitamin biosynthesis, carbohydrate utilization, short chain fatty acid levels and more (58, 60), making it likely that metabolomic measurements would vary between infants and those at later maturation stages. Consistent with this, we observed age-related variation across multiple metabolite classes, including peptides, carboxylic acids, lipids, amino acids, all of which may be influenced by shifts in gut microbial composition.

The analysis described here has several limitations. Due to its exploratory nature, the sample size was small and the analysis may have been underpowered to detect differences between the age cohorts, particularly when the magnitude of the differences was modest. This limitation was especially relevant for the post-pubertal age group and may have contributed to the smaller number of metabolite concentration differences observed relative to the pre-teen group after Bonferroni correction. In addition, no biochemical or clinical markers were available to confirm maturation stage or pubertal status; therefore, intra-group developmental heterogeneity likely existed. The cohort was also predominantly male, which, together with the small size of the post-pubertal group, precluded meaningful analyses of sex-related differences. Nonetheless, despite the relatively small cohort size, this analysis identified age-related trends in more than 20% of measured metabolites within the plasma metabolome, many of which have not been previously reported. These findings provide a rationale for future studies aimed at more comprehensively characterizing age- and sex-related variation in plasma metabolite concentrations across the pediatric age spectrum.

The findings presented here suggest that there is age-related heterogeneity in the pediatric plasma metabolome, and that the use of a single “pediatric” normal range for certain metabolites is inappropriate. Indeed, age-related changes from body size, gut microbiome and sexual maturation intuitively suggest that associated changes in the metabolome should be expected. These data signal the need for more detailed characterization of the plasma metabolome across the pediatric age spectrum, with additional accounting for biological sex and pubertal stage. Such normative reference ranges will be essential to improve the accuracy of metabolomic analyses that include pediatric participants.

## Supporting Information

The following files are available free of charge.

Supplementary Figures (PDF)

## Data Availability

This study is available at the NIH Common Fund’s National Metabolomics Data Repository (NMDR) website, the Metabolomics Workbench (61), https://www.metabolomicsworkbench.org where it has been assigned Study ID ST004827 and Project ID PR003073. The data can be accessed directly via its Project DOI: http://dx.doi.org/10.21228/M8G27K

## Author Contributions

The manuscript was written through contributions of all authors. All authors have given approval to the final version of the manuscript. Amy Thachil participated in research design, data analysis and writing of the paper. Rupasri Mandal participated in performance of the research and writing of the paper. David Wishart participated in research design, performance of the research and writing of the paper. Tom Blydt-Hansen participated in research design, data analysis and writing of the paper. The authors declare no conflicts of interest.

## Funding Sources

No funding to disclose.

## Abbreviations

ANOVA: analysis of variance
CALIPER: Canadian Laboratory Initiative on Pediatric Reference Intervals
Cers: ceramides
CEs: cholesteryl esters
DI-MS: direct injection mass spectrometry
DGs: diglycerides
ENT: ear, nose and throat
HMDB: Human Metabolome Database
LC/MS-MS: liquid chromatography mass spectrometry
LOD: limit of detection
LPC: lysophosphatidylcholines
PCA: principal component analysis
PCs: phosphatidylcholines
SMs: sphingomyelins
TGs: triglycerides

## Notes

https://metabolomicsworkbench.org/data/DRCCMetadata.php?Mode=Study&StudyID=ST004827

### Competing Interest Statement

The authors have declared no competing interest.

