## Supplemental Figure 1 for "Age-related differences in metabolome composition in a healthy pediatric population: A pilot analysis"

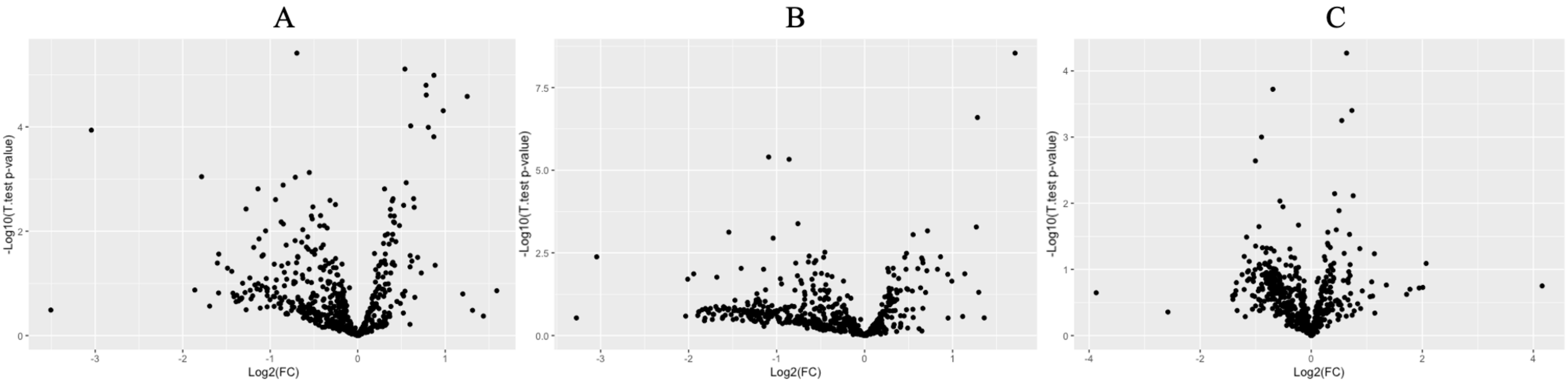


**Figure S1. Fold change differences in metabolite concentrations between developmental stages.** Overall, comparisons between the pre-teen and infant age groups yielded the greatest number of differences followed by the post-pubertal versus infant and post-pubertal versus pre-teen groups. T-test results demonstrated that 99 metabolites were different between the pre-teen and infant age groups, 73 metabolites demonstrated a significant concentration difference between the post-pubertal and infant age groups, and 26 metabolites were different between the post-pubertal and pre-teen groups ($p<0.05$). **A)** Pre-teen/infant metabolite differences. Fold changes were calculated in the following direction: pre-teen/infant. 11 metabolites demonstrate a fold change <0.5 between the two groups and 1 metabolite demonstrates a fold change >2 ($p<0.05$). **B)** Post-puberty/infant metabolite differences. Fold changes were calculated in the following direction: post-puberty/infant. 10 metabolites demonstrate a fold change <0.5 between the two groups and 5 metabolites demonstrate a fold change >2 ($p<0.05$). **C)** Post-puberty/pre-teen metabolite differences. Fold changes were calculated in the following direction: post-puberty/pre-teen. 3 metabolites demonstrate a fold change <0.5 between the two groups and no metabolites demonstrate a fold change >2 ($p<0.05$).
